# How and why to quantify pairwise pleiotropy and genotype-by-environment interactions

**DOI:** 10.64898/2026.01.09.698568

**Authors:** Thomas James Ellis

**Affiliations:** Uppsala University, Sweden; Gregor Mendel Institute of Molecular Plant Sciences, Vienna Austria

## Abstract

Pleiotropy is when a single locus affects two or more traits. The magnitude and direction of pleiotropy can constrain or faciliate phenotypic evolution. Investigations of pleiotropy have typically relied on null-hypothesis tests to classify cases into discrete categories based on the direction of effects. This discrete approach ignores the quantitative nature of pleiotropy, and systematically underestimates pleiotropic interactions. I describe a simple method to quantify the direction and magnitude of pleiotropic effects to alleviate these issues for pairs of traits. I illustrate how genotype-by-environment interactions can be viewed as a special case of pleiotropy and described in the same way. I provide an R package, `psiotropy`, to apply these methods.

## 1. Introduction

Pleiotropy is when a single locus affects two or more traits. It is of particular interest in evolution because it can either facilititate evolutionary change when multiple traits are selected in parallel, or constrain it when traits trade-off with one another. Pleiotropy is thought to play an important role in trade-offs between life history traits such as longevity and reproduction, in adaptation to contrasting environments, and in the degree to which phenotypic complexity constrains evolution (Williams, 1966, 1957; Stearns, 1992; Kawecki and Ebert, 2004; Wagner et al., 2008). One fruitful line of research has been to characterise pleiotropy by measuring its effects on correlations between phenotypes (Stearns, 1992). With the more recent advent of molecular and statistical techniques to identify the genetic basis of traits we have begun to address the challenges involved in understanding the pleiotropic mechanisms acting at individual loci (Mitchell-Olds et al., 2007; Wagner and Zhang, 2011; Wadgymar et al., 2017; Jee et al., 2025).

Key to understanding these evolutionary outcomes is the direction or mechanism of pleiotropy. Three mechanisms are typically invoked to describe pleiotropic effects of alleles on pairs of traits (Kawecki and Ebert, 2004; Mitchell-Olds et al., 2007; Hall et al., 2010). First, negative (or antagonistic) pleiotropy occurs when an allele is associated with an increase in one trait, but a decrease in a second trait. A second ‘non-pleiotropic’ case occurs when an allele is associated with a change in a single trait, but no change in the second. When the traits in question are fitness in two environments, the latter case is often referred to as ‘conditional neutrality’ (Mee and Yeaman, 2019). Finally, positive pleiotropy occurs when an allele is associated with either an increase or a decrease in both traits. These mechanisms have obvious parallels to negative, zero, and positive genetic correlations between overall phenotypes. The strong or more common each of these directions of pleiotropy are, the more we would expect overall genetic correlations to be. The frequency of each pleiotropic mechanism is therefore central to determining the relationships between traits, and how they constrain one another.

To understand trade-offs it is essential to be able to measure them. For example, relationships between traits can be measured as genetic correlations (Falconer and Mackay, 1996; Stearns, 1992). The resulting correlation coefficient provides a quantitative and intuitive measure of the direction and magnitude of the relationship, which can be easily compared across studies. However, this relies on having a sample of many genetically diverse individuals on which to measure phenotypes. To investigate the pleiotropic effects of individual loci we can only measure the effect of one allele against the other on each trait in question. A typical approach is to perform a statistical test on the effect on each trait, and classify loci as affecting neither, one or both traits. However, as I outline in detail below, this classification process ignores the fundamentally continuous nature of the problem and leads to a statistical bias that systematically underestimates the amount of pleiotropy in the system (Hill and Zhang, 2012). Just as classical correlation coefficients measure both the direction and the strength of a relationship between traits, what we need is a framework to quantify the direction and strength of pleiotropic effects.

In the rest of this paper I highlight three key reasons why current approaches to measuring pleiotropy are flawed, and use these to motivate a quantitative framework for measuring pleiotropy between pairs of traits. I then illustrate how genotype-by-environment interactions can be viewed as between-environment pleiotropy, and described in the same way. I provide an R package psiotropy to implement the methods described in this paper. It can be accessed from GitHub at https://github.com/ellisztamas/psiotropy.

## 2 Null-hypothesis tests bias conclusions

I consider the general case of two genotypes, A and B, that have been measured for two traits. Usually, A and B would be different genotypes at a single locus, but what follows is equally applicable to a comparison of genetic or breeding values of two lines that differ genome wide. The two genotypes differ in trait 1 by effect size *x* and in trait 2 by effect size *y*. The best way to define effect sizes will depend on the organism and study design, but would typically be the difference in phenotype between A and B. In other cases, such as when the traits are measures of fitness or its components, the (log) ratio of phenotypes might make more sense.

The next step typically involves some kind of statistical test of the null hypotheses that *x* and/or *y* are zero. A common approach is to perform two univariate tests for each trait separately, often accompanied by a discussion of the relative direction of the effect on each trait (e.g. Wagner et al., 2008; Hall et al., 2010; Anderson et al., 2011; Ågren et al., 2013; Ellis et al., 2021). Variations on the testing scheme exist, including multivariate extensions; see Porter and O’Reilly (2017) for a review. Loci that are significantly associated with both loci are classified as positively or negatively pleiotropic (the green and red points in figure 1A), those associated with only one trait as showing zero pleiotropy (blue points in figure 1A), and those affecting neither trait are ignored (black points in figure 1A). Another approach is to identify clusters of multiple covarying traits for further study (reviewed in Jee et al., 2025), although in these cases the mechanism of pleiotropy is usually not the focus, and I do not consider them further. For null-hypothesis-testing approaches the goal is to classify each locus as affecting one, both, or neither trait.

**Figure 1:**
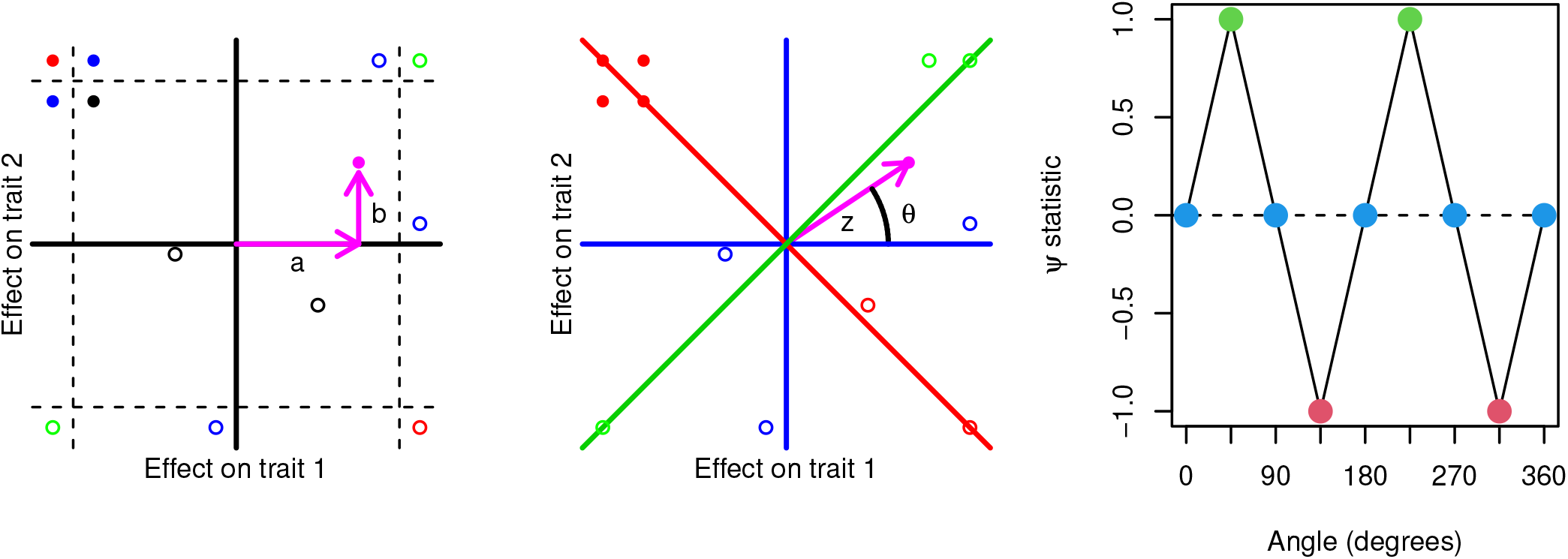
Two ways to measure pleiotropy. Each point shows the effect of a hypothetical locus on traits 1 and 2. Colours indicate pleiotropic mechanism: red, green and blue points indicate loci showing negative, positive and zero pleiotropy; black points show no significant association with either trait. (A) Null-hypothesis tests of allelic effects on each trait. A reference allele at each locus has effects *x* and *y* on traits 1 and 2 respectively, and the statistical tests are performed along the x- and y-axes. The grey bars indicate the boundary of statistical significance. Each locus is categorised as affecting zero, one or both traits based on which side of the bars they are located. (B) The same loci, but represented as vectors with angles (*θ*) and lengths (of magnitude *z* ) from the origin. (C) Function *ψ* relates angles to pleiotropic mechanisms. Three special cases when *ψ* is -1, 0 or 1 correspond to antagonistic pleiotropy (red), zero pleiotropy (blue) and positive pleiotropy (green) respectively. Values between these points measure everything in between.

There are three shortcomings with this approach. First, by classifying effects based on significance thresholds we ignore useful quantitative information about pleiotropic mechanisms. For example, if we classify the loci in figure 1A based on how many traits they each affect we would infer that the red loci show negative pleiotropy, the blue loci no pleiotropy, and we would ignore the black loci as non-significant. In fact it is clear that the actual effects are often much closer to those of loci classified as having different pleiotropic mechanisms than they are to loci classified as having the same pleiotropic mechanism. As an example, look at the solid points in figure 1A; the effects of these loci are all broadly associated with a decrease in trait 1 and an increase in trait 2. However, they happen to fall on either side of the significance thresholds, and are classified as having distinct pleiotropic mechanisms. By classifying pleiotropic mechanisms into discrete categories based on statistical significance at individual traits we ignore quantitative information about the strength and mechanism of pleiotropy. In this way we risk allowing statistical significance to obscure biological meaning.

Second, identifying pleiotropy based on how many significant tests we find requires that we detect two or more statistically significant associations. However, it will always be more difficult to find two significant associations than one (Wagner and Zhang, 2011; Hill and Zhang, 2012). Using a standard significance threshold of p=0.05, the a priori expectation of identifying an association with a single trait is 0.05. The expectation of finding two significant associations is *p*^2^ = 0.00625, or twenty-fold less likely. Higher-order pleiotropic interactions become exponentially more difficult to find. For a visual intuition for this, observe that loci classified as pleiotropic in figure 1A must fall into the corners of the diagram. These are the very regions that are furthest from the centre, meaning that effects have to especially strong to be classified as pleiotropic. This means that inference of the direction of pleiotropy is confounded with the magnitude of the the effect. This leads to a statistical bias that systematically underestimates the amount of pleiotropy in the system (Hill and Zhang, 2012).

Consistent with this, early estimates of the extent of pleiotropy based on hundreds or thousands of samples, usually in model organisms like mice and yeast, found few examples of statistical associations with multiple traits, and concluded that pleiotropy is limited (Wagner and Zhang, 2011). In contrast, more recent work in humans with hundreds of thousands of samples have found that a large fraction of the genome shows pleiotropy (e.g. Boyle et al., 2017; Watanabe et al., 2019). Unless humans show unusually high levels of pleiotropy, this discrepancy must be due to differences in sample size, and hence statistical power. It is unlikely that such sample sizes will become available outside humans soon, if at all, so we require a way to describe pleiotropy which minimises the bias due to sample sizes.

Third, use of significance thresholds biases the stength of effect sizes deemed to be important. It is well known that unless sample sizes are large, conditioning on significant associations tends to overestimate effect sizes (Beavis, 1998; Xu, 2003). This also means we filter out any loci deemed to be ‘non-significant’. The extent to which these matter will depend on the biological question, but quantitative traits are generally expected to have a substantial contribution from loci with weak to moderate effects, but which can have a substantial contribution to genetic variance in aggregate Fisher (1930). Rather than remove them, a stronger approach would be to quantify how much weak effects contribute to overall trade-offs.

In summary, by relying on null-hypothesis testing to infer mechanisms of pleiotropy we (1) discretise what is really a continuous phenomenon, (2) underestimate the amount of pleiotropy, and (3) ignore the contribution of weak effects. To alleviate this, we would prefer to (1) quantify rather than classify the direction and magnitude of pleiotropy; (2) focus on the estimation and uncertainty of those measures, in a way that decouples one from the other; and (3) uses as much of the data as possible.

## 3 Quantifying trade-offs

### 3.1 Effects as angles and magnitudes

I propose a simple approach to address the goals identified in the previous section. Rather than define pleiotropic effects along the axes of traits 1 and 2 (figure 1A) we can instead describe effects as angles and vector lengths. Any pair of values for *x* and *y* can be described without loss of information as angle *θ* and magnitude 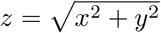 (figure 1B).

Angle *θ* contains information about the direction of pleiotropy. When *θ* points along the red, blue and green axes in figure 1B these correspond to loci showing negative, zero, and positive pleiotropy respectively. *θ* can be turned into a more useful statistic by taking function *ψ* (figure 1C) This function is cyclical and hence must be defined piecewise:

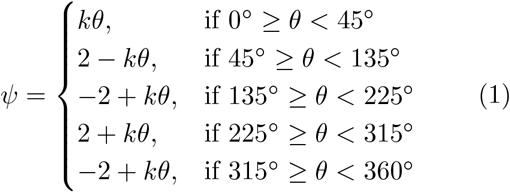

where 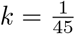. This resembles a linearised sine wave, and is likely to be clearer as a figure (figure 1C) than an equation. Values of *ψ* of -1, 0, and 1 correspond to the special cases of symmetrical negative, zero and positive pleiotropy (red, blue and green in figure 1B and C). However, since *ψ* is quantitative, all intermediate values between these cases are possible. In this way, *ψ* reflects an intuitive quantitative measure of the pleiotropic mechanism.

The length *z* of the vector reflects the overall magnitude of the effect. Just as larger values of *x* and *y* reflect stronger effects on individual traits, values of *z* which are further from the origin reflect a stronger effect overall. A key difference between these numbers is that *x* and *y* measure effects along specific axes, but *z* but is agnostic about the direction of the effect. This decouples estimation of the direction from the magnitude of the effect. This also means that estimates of uncertainty can be assessed on one axis only, negating the need to test multiple traits at once (see below).

Two conditions must be met for *ψ* and *z* to be meaningful. First, it is essential that *x* and *y* are on the same scale so they can be compared. For quantitative phenotypes this can be acheived by standardising phenotypes by their mean and standard deviation, and estimating effect sizes on the scaled values (Schielzeth, 2010). When effect sizes are ratios, one can use the log of the ratio. Second, it is essential that *x* and *y* be defined so that the choice of reference allele is arbitrary. This means that if the genotype labels were swapped, effect sizes would simply swap sign. In bivariate space, this would cause the vector to be reflected across the origin, which will return the same value of *ψ* as if the sign had not been flipped (figure 1B). This is important because the definition of ‘reference’ and ‘alternative’ allele is typically arbitrary.

### 3.2 Statistical robustness

#### 3.2.1 Sampling distributions of pleiotropy statistics

In addition to point estimates of *ψ* and *z* one would typically wish to estimate the uncertainty of these statistics. Two issues should be borne in mind here. First, if we were to compare vectors of different magnitudes of *z*, we would expect shorter vectors to have higher uncertainty in *ψ*, even if the standard errors of *x* and *y* remain constant. Figure 2A shows values drawn from three distributions with equal standard deviations, but increasingly far from from the origin. The spread of the three distributions in *x* and *y* is very similar, but the range of angles with respect the origin increases between the ochre and green points, reflected in wider confidence intervals on *ψ*. As *z* approaches zero (purple), the overlap becomes complete, and the distribution spans all directions from the origin, and *ψ* ranges from -1 to 1. The stronger the allelic effects the more robust are estimates of the direction of pleiotropy.

**Figure 2:**
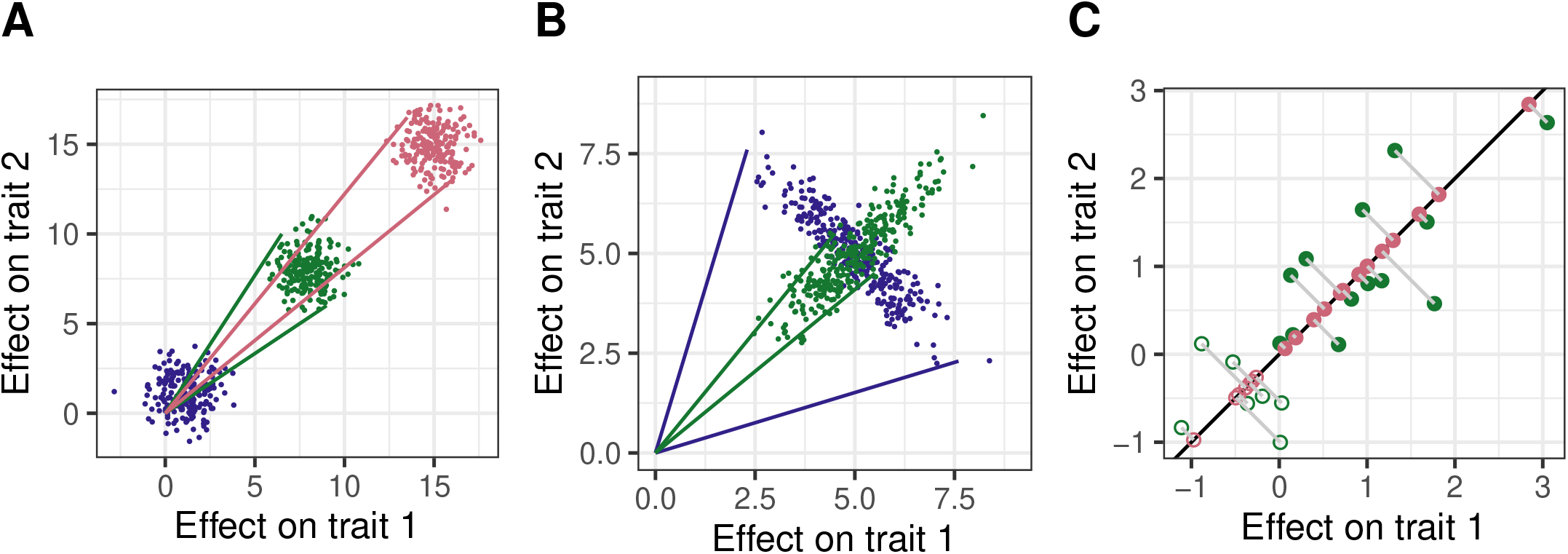
Measuring uncertainty in pleiotropy statistics. (A) Sampling distribution of 200 random deviates from bivariate normal distributions at increasing distances from the origin. Solid lines indicate the approximate confidence intervals of the range of angles in each distribution; blue points reflect angles in all directions. (B) Sampling distribution of 300 random deviates from bivariate normal distributions with equal means and standard deviations, but covariances of opposite sign. (C) Illustration of *z*^∗^ and *q*. Green points show 20 random deviates from a bivariate normal distribution. Ochre points show the project of those points onto vector *v*. Shaded and filled points indicate with projected values are greater or less than zero along *v*; in this case *q* = 7*/*20 = 0.35.

Second, traits of interest are often genetically correlated, especially if there is pleiotropy. This in turn causes the sampling distributions of *x* and *y* to be correlated. The direction of this correlation can have a major impact on uncertainity in *ψ*. Figure 2B shows values drawn from two distributions with equal means and standard deviations (and hence equal expected values of *ψ* and *z*), but opposite signs of the correlations between errors. When the directions of *ψ* and the correlation of sampling distributions are closely aligned we expect less uncertainty in *ψ* (green). When the directions are opposite, we expect greater uncertainty in *ψ* (purple). By extension, if we were to incorrectly assume the correlation were zero we would estimate the uncertainty in *ψ* incorrectly. As such, care must be taken that any estimate of uncertainty accounts for the correlation between the sampling distributions of *x* and *y*.

With these points in mind, a simple way to estimate uncertainity in pleiotropy statistics is to generate a sample of estimates *x*^∗^ = (*x*_1_, …, *x*_*n*_) and *y*^∗^ = (*y*_1_, …, *y*_*n*_) around the point estimates *x* and *y*, and calculate 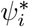 and 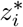 for each pair of values. It is then straightforward to calculate confidence intervals from the quantiles of those distributions. This sample can easily be generated by non-parametric bootstrapping, which when done correctly should preserve the covariance structure between *x* and *y* (Efron, 1982). Other approaches would be parametric bootstrapping or Bayesian regression, as long as care is taken to model the covariance. Samples of estimates of *x* and *y* thus provide a simple and robust way to estimate uncertainty in pleiotropy statistics.

#### 3.2.2 Quantifying statistical robustness

As mentioned above, the closer *z* is to zero, the more the distribution of *x*^∗^ and *y*^∗^ overlap zero, and the greater the uncertainty in *ψ*. Thus, the degree of overlap reflects a quantitative estimate of the robustness of estimates of *ψ*. This is directly analogous to estimating a confidence interval around a one-dimensional measurement, and asking the extent to which that interval overlaps zero. Since *x* and *y* vary in two dimensions, we wish to simplify the problem and quantify the extent to which confidence intervals in *z* overlap the origin. If the observed vector the observed vector (*x, y*) is *v*, we can project each vector 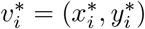 of sampled values in onto *v*:

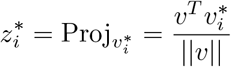

These projected values now vary only in one dimension along *v* (figure 2C). For that reason they are interpretable as magnitudes along a vector, and we can call them 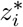. We then calculate value *q* of (twice) the proportion of 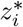 less than zero (i.e. that fall on the opposite side of the origin to *v*). When pairs of resampled points do not overlap zero, *q* = 0. As *z* approches zero and overlap become complete, *q* approaches one. *q* is thus a convenient summary statistic for the extent to which a pleiotropic vector overlaps zero.

*q* is particularly useful when there are lots of loci to summarise, and it is impractical to visually inspect each one. One approach would be to treat *q* as analagous to a *p*-value, and filter out loci that fall below some threshold of significance, such *q* = 0.05. This would not introduce the same bias caused by requiring significant tests for every trait described above, because the such a test would be independent of the direction of pleiotropy. However, it would still impose a threshold that is fundamentally arbitrary, which risks obscuring interesting but weak effects, and takes us back to classifying rather than quantifying pleiotropic effects. Alternatively, one could use 1− *q* as a quantitative score to weight the effects of particular loci in downstream analyses. This does not introduce arbitrary thresholds, but does use the full information about uncertainty in the underlying point estimates.

## 4 Genotype-by-environment interactions

The previous sections focussed on pleiotropy between multiple traits measured in the same individuals. A special case of pleiotropy arises when a single trait is measured in multiple environments (Falconer, 1952). I focus on an important example of between-environment pleiotropy that arises in the study of local adaptation; theory predicts that locally-adapted genotypes should have higher fitness in their native environment, and reduced fitness in other environments (Kawecki and Ebert, 2004). Nevertheless, this applies to genotype-by-environment interactions in general.

The typical approach to identify and interpret between-environment pleiotropy is somewhat different from between-trait pleiotropy. Environmental interactions lend themselves well to ANOVA and related linear models that include main effects of genotype and environment, and a genotype-by-environment interaction effect. A significant interaction term indicates between environment pleiotropy. The meaning of the interaction is usually interpreted in the context of a reaction-norm plot (Fig 3A-E). Crossing reaction norms indicate ‘antagonsistic pleiotropy’, where different genotypes are favoured in each environment, and hence that there is a trade-off between environments. An effect in one environment but not the other indicates ‘conditional neutrality’; one genotype is fitter in one environment, but there is no fitness difference in the other (Mee and Yeaman, 2019). Parallel reaction norms indicates that one genotype shows ‘global superiority’, reflected in higher fitness in both environments. The combination of linear models and reaction-norm plots has been a very informative tool for investigation between environment pleiotropy.

**Figure 3:**
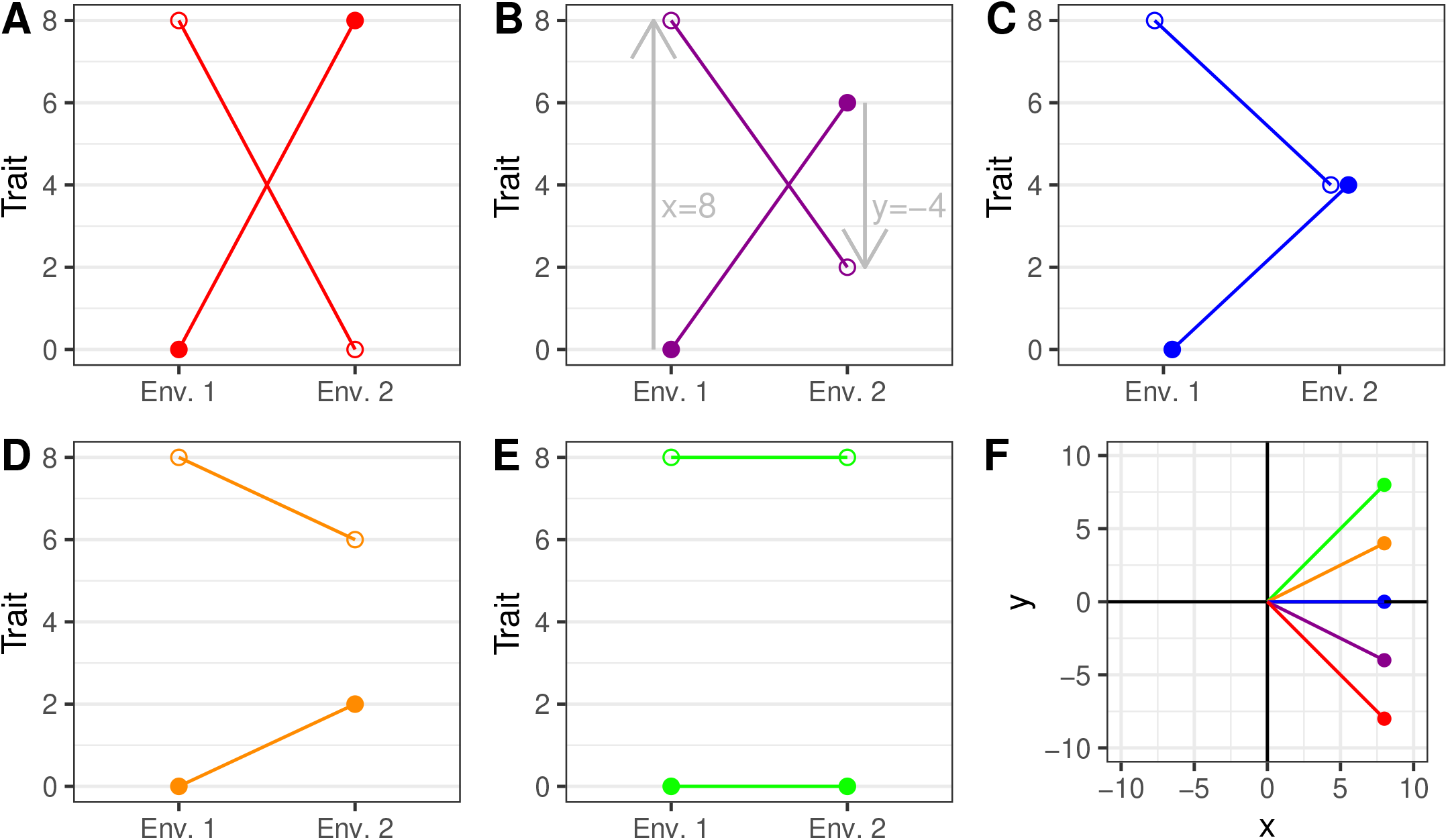
Genotype-by-environment interactions as between-environment pleiotropy. Plots show hypothetical trait values for genotypes A (open circles) and B (closed circles) in two environments. A-E show reaction-norm plots for a continuum of scenarios ranging from (A) a symmetric genotype-by-environment interaction (C) a genotypic effect in only one environment, to (E) a genotype-effect only. B and D are intermediate scenarios. Grey arrows in B illustrate one way to calculate effect sizes *x* and *y* as the differences in phenotype between genotypes A and B in each environment. F illustrates effect sizes for A-E plotted as vectors in two dimensions. Colours correspond to those in A-E.

Nevertheless, the approach has several shortcomings. First, the process implicitly classifies interactions into discrete cases of anatagonistic pleiotropy, conditional neutrality and global superiority. Just as for between-trait pleiotropy, there is really a continuum between these cases, and we would like a way to quanitify that continuum. Second, it is not straightforward to interpret patterns of pleiotropy from the main and interaction statistics of a linear model directly, without also visually inspecting the reaction-norm plot. This is fine when there are relatively few interactions to check, but quickly becomes impractical for large numbers of interactions, such as pleiotropy at loci across the genome. Finally, using null-hypothesis tests to guide inference again introduces confounding between the direction and magnitude of pleiotopy. The null hypothesis test for a genotype-by-environment interaction is that *x* = *y*, and hence that *ψ* = 1. Conditioning on loci that show a significant genotype-by-environment interaction thus biases inference towards finding loci showing global superiority. As for between-trait pleiotropy, quantifying between environment pleiotropy would be a useful way to augment existing approaches.

Fortunately we can quantify the direction and magnitude of between-environment pleiotropy just as easily as between-trait pleiotropy. In general, cases of antagonistic pleiotropy should show *ψ* ∼ − 1, cases of conditional neutrality should show *ψ*∼ 0, and cases of global superiority should show *ψ*∼ 1. For ease of presentation, we might define *x* and *y* in the scenarios depicted in figure 3A-E as simply the difference in phenotype between genotypes A and B in each environment (see the grey arrows in figure 3B). In these scenarios *x* = 8 in all cases, and *y* is -8, -4, 0, 4 and 8 respectively. These values can easiy be expressed as vectors (figure 3F) and used to calcuate pleiotropy statistics, giving values of *ψ* of -1, -0.59, 0, 0.59 and 1 respectively. In this way *ψ* can be viewed as a measure of the extent to which reaction norms are (or are not) parallel, which is a biologically meaningful measure of the extent to which traits differ between environments.

## 5 Discussion

Standard approaches to measuring pleiotropy have treated the problem as a square or rectangle, whose widths and heights are defined by individual traits (figure 1A). In the square framework, the mechanism of pleiotropy is classified into discrete states, but that classification is confounded with the strength of the effect. I argue that pleiotropy is better viewed as a circle, with angle *θ*, radius *z*, and pleiotropic mechanism *ψ* (figure 1B). In the circular framework, *ψ* and *z* provide quantitative measures of the direction and strength of pleiotropy that obviate the need to classify, and decouple direction and magnitude. Ignoring the contradiction between the circle and the square is like trying to push a round peg through a square hole; the only way to do it is to chisel off parts of the peg, and in doing so lose something fundamental about the circle. Treating pleiotropy as a circle allows us to gain a less biased picture of the nature of pleiotropy.

Our method relies on good estimates of effect sizes *x* and *y*. I have highlighted that the best way to define these will depend on the biological question at hand, and I encourage users to invest time into thinking about how best to do this. For example, estimates are likely to be different if relevant experimental or population confounders are accounted for, such as temporal or spatial batch effects (Paaby and Rockman, 2013; Solovieff et al., 2013). Particular concerns for the investigation of pleiotropy include linkage between multiple loci affecting individual traits, and genetic confounding due to population structure between loci affecting separate traits. Furthermore, assumptions about the causal relationship between traits should affect estimates of the correlation between genotype and each phenotype. A persistent issue is the extent to which pleiotropy is ‘horizontal’ (the gene has independent effects on both traits) or ‘mediating’ (the gene affects one trait, which then affects the second). These issues are of course not limited to the method presented here; null-hypothesis tests also rely on *x* and *y*. Fortunately, *ψ* can be calculated with little computational burden, making it easy to compare results based on different estimates. Quantifying pleiotropy thus does not solve issues of confounding, but my hope is that it makes it easier to interpret.

There is a close analogy between *ψ* and traditional correlation coefficients. Just as correlation coefficients of -1, 0 and 1 correspond to negative, zero and positive correlations, values of *ψ* of -1, 0 and 1 correspond to negative, zero and positive pleiotropy. Indeed, an interesting avenue of future research would be to quantify the extent to which the direction of genotypic correlations at the phenotypic level is refelected in the distribution of values of *ψ* across the genome without the bias caused by null hypothesis tests. *ψ* is thus very similar to estimating a correlation, except that correlations are between pairs of vectors numbers and *ψ* is between pairs of individual point estimates. As such, it is more broadly applicable beyond questions of pleiotropy whenever it is only possible to estimate a single pair of point estimates.

The method outlined here describes pairs of traits. It is not immediately clear how to directly extend the statistics *ψ* and *z* to three or more traits. That said, I note that it also not clear how to extend traditional correlation statistics to three or more traits, but this has not prevented them from being useful. One potential avenue of research would be to estimate *ψ* for multiple pairs of traits, and to use those values as edges in a ‘pleiotropic network’. This is analagous to the idea of a gene co-expression network, where edges are built on correlation coefficients. The pleiotropic network could then be investigated using the tools of network analysis.

In conclusion, I have presented a simple approach to describe the direction and magnitude of pleiotropic effects on pairs of traits. I also provide an R package to allow users to apply the method to their own data. I hope this provides the basis for future work investigating the quantitative nature of pleiotropy.

## 6 Acknowledgements

I thank Jon Ågren and Pascal Milesi for constructive feedback on previous versions of this manuscript.

